# Unveiling Candidate Drivers in Cancer Progression Using Somatic-IV Analysis

**DOI:** 10.1101/2025.02.20.638369

**Authors:** Wei Zhang, Saba Ghaffari, Andrew Browne, Carlos Macintosh, Xiaoyu Lu, Ha Dang, Gonzalo Lopez, Kai Wang

**Affiliations:** Bristol Myers Squibb; University of Illinois Urbana-Champaign; Independent Professional

## Abstract

**Motivation:** Identifying causal drivers of cancer progression is crucial for developing effective anti-cancer therapies. However, disentangling causality from correlation remains challenging due to confounding factors within the complex genetic and transcriptomic landscapes of cancer.

**Results:** To address this challenge, we introduce the Somatic Instrumental Variable analysis (Somatic-IV), which integrates genetic, transcriptomic, and clinical outcome data to identify candidate driver genes likely playing a causal role in disease progression. Somatic-IV estimates genetic-exposure and genetic-outcome associations, utilizing MR-Egger regression to estimate bias-reduced causal effects. We assessed the statistical properties and performance of Somatic-IV through a simulation study and demonstrated its utility in pancreatic adenocarcinoma (PDAC) and colorectal cancer (CRC). Our analysis identified ZBED2 and CSNK2A2 as likely drivers of PDAC and CRC cancer progression, respectively. Somatic-IV is a novel approach for generating target hypotheses and guiding the development of effective therapies for cancer patients.

**Availability and implementation:** The Somatic-IV package is available at: https://github.com/wzhang1984/Somatic_IV_package

## 1. Introduction

Disease progression and treatment resistance remain major challenges in cancer research and patient care. Identifying the key dependencies and mechanisms of cancer progression and treatment resistance is crucial to developing effective therapies and improving patient outcomes. However, the heterogeneity of tumors in both genetic and transcriptomics landscapes complicates the identification of cancer drivers, posing a significant challenge to this goal.

Genetically, while a few genes carry frequent mutations or copy number events, most somatic alterations occur at low frequency. Many desirable cancer drivers with recurrent somatic alterations have been difficult to drug, such as the MYC oncogenes (Dang, et al., 2017). Although preclinical data suggest potential efficacy for targeting some rare mutations (Jiang, et al., 2013), the low prevalence of these alterations has limited the interest in exploring therapies against these genetic drivers. As a result, developing a targeted therapy that benefits a large patient population has been challenging. However, recent studies have shown that upstream somatic alterations converge on a set of downstream drivers that integrate these signals and govern the transcriptional program of cancer cells (Bradner, et al., 2017; Paull, et al., 2021). Targeting such transcriptional drivers holds promise for encompassing a broader spectrum of patients.

Transcriptomics-based subtypes have been extensively studied in many types of cancer, based on bulk tissue or single-cell RNA sequencing, with the aim of homogenizing the patient population and combating genetic heterogeneity. Despite these efforts, transcriptomics-based patient stratification rarely informs clinical decisions or treatment development. The associations between somatic alterations and transcriptionally defined subtypes are often weak, posing a challenge in identifying transcriptional drivers that mediate the impact of upstream genetic alterations to downstream transcriptional states.

One promising approach to address these challenges is to study the influence of genetic variants on clinical outcomes through the modulation of key genes or mechanisms. In human genetics, powerful strategies like Mendelian Randomization (MR) (Bowden, et al., 2015) and transcriptome-wide association study (TWAS) (Gamazon, et al., 2015) have been developed based on this principle. These causal inference methods use genetic variants as instrumental variables (IVs) to estimate the potential causal effect of the genetically regulated portion of an exposure variable (e.g., gene expression) on an outcome of interest in observational studies. However, this type of strategy has not yet been extended to the studies of cancer somatic alterations and clinical outcomes.

In this paper, we introduce a computational framework for identifying mechanisms of cancer progression by an IV-based causal inference approach, building on the concept of MR and extending it by utilizing cancer somatic alteration events as IVs. Our framework aims to identify likely drivers or driving mechanisms that mediate the impact of somatic alterations on disease progression in cancer patients. We present the applications of our computational framework in two disease indications: pancreatic adenocarcinoma (PDAC) and colorectal cancer (CRC). Our computational framework offers a novel approach to generate target hypotheses, which we hope could facilitate the development of novel and effective therapies for cancer patients.

## 2. Material and Methods

In the upcoming sections, we first provide a brief overview of the MR method. We then introduce our novel framework, which extends the MR analysis to identify likely drivers and driving mechanisms of cancer progression.

### 2.1. Causal inference using Mendelian Randomization

The Mendelian Randomization (MR) method was originally developed in epidemiology to establish causal relationships between an exposure and an outcome (e.g., the causal link between blood pressure and stroke). Unlike randomized controlled trials, which are often costly and impractical, MR utilizes genetic variants (e.g. SNPs) as instrumental variables (IVs) that mimic the randomization process. This method provides a powerful way to infer causality from observational data by making the following key assumptions: (1) IVs are associated with exposure, (2) IVs are independent of confounders, and (3) IVs are associated with the outcome only through the exposure. When these assumptions are met, MR can reduce both reverse causality and confounding (Haycock, et al., 2016). In practice, MR can be implemented using two common approaches: two-stage least squares (TSLS) (Angrist and Imbens, 1995) or MR-Egger regression (Bowden, et al., 2015).

MR involves analyzing data from a study on *N* patients. For each patient *i*, the study collects data on *J* genetic variants (*G*_*i*1_, *G*_*i*2_, … *G*_*iJ*_), an exposure variable (*X*_*i*_, e.g., gene expression, protein activity), an outcome (*Y*_*i*_), and considers known and unknown confounders on both the exposure 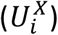 and the outcome 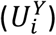. The exposure is modeled as a linear combination of genetic variants, the confounders, and an independent error term 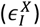 (**Eq. 1**), where *γ*_*j*_ represents the effect size of genetic variant *j* on the exposure. The outcome is then modeled as a linear combination of the genetic variants, the exposure, the confounders, and an independent error term 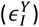 (**Eq. 2**), where *β* represents the causal effect of the exposure on the outcome, and *a*_*j*_ represents the direct effect of variant *j* on the outcome without being mediated by the exposure.

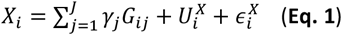

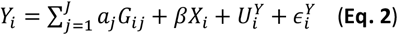

### 2.2. Validity of using cancer somatic alterations as IVs

To apply the MR technique to identify oncogenesis drivers that contribute to tumor progression, we investigate the potential use of cancer somatic alterations, including somatic mutations and copy number alterations (CNAs), as IVs. Valid IVs must satisfy 3 key assumptions outlined in Section 2.1. To address assumption 1, we introduce a network-regularized Lasso technique that ensures both statistical and biological associations between somatic alteration IVs and the candidate drivers of interest (i.e. the exposure variable), as detailed in Section 2.3.3. To account for assumption 3, we employ the MR-Egger technique to correct for bias due to violations of this assumption (Section 2.3.2).

However, satisfying assumption 2, which requires somatic alternations (i.e. IVs) to be independent of all confounding factors, presents a challenge. While the central dogma suggests that somatic mutations should not be influenced by confounders such as downstream transcriptional states and metabolic pathways, we acknowledge that confounders may still violate assumption 2 in two main ways. Firstly, a confounder (e.g., age) may increase the likelihood of certain somatic mutations occurring and also lead to adverse effects that result in unfavorable patient outcomes. Secondly, a confounder, such as targeted therapies, may selectively impact patient subpopulations with specific somatic alterations, thereby influencing patient outcomes positively. Consequently, careful patient sample and IV selection is crucial. For IV selection, we empirically tested all available demographic and clinicopathological variables (i.e. age, gender, race, tumor stage, tumor grade) in the patient datasets and excluded somatic alterations associated with them (Sections 3.2.1 and 3.3.1). Regarding patient sample selection, we excluded patient subgroups with known somatic alternations associated with targeted therapies (Section 3.3).

### 2.3. Causal inference of cancer progression drivers using modified Mendelian Randomization

We introduce a novel framework called Somatic Instrumental Variable approach (Somatic-IV) to estimate the causal effects of molecular drivers on cancer progression (**Fig. 1**). Our framework investigates the activity of a candidate patient state driver (PSD) as the exposure of interest, and utilizes two modifications to extend the traditional MR approach. First, we model somatically acquired mutations and copy number alterations (CNAs) as IVs instead of inherited SNPs. Second, our framework uses a censored time-to-event outcome, such as overall survival (OS) or progression-free interval (PFI) of cancer patients. To analyze patient survival as outcome, we adapt the MR approach using the Aalen additive hazard model (Aalen, 1989; Tchetgen Tchetgen, et al., 2015). Specifically, for each patient *i*, the hazard function of this outcome, *T*, evaluated at time *t*, can be conditioned on the PSD’s activity *X*_*i*_, somatic alterations *G*_*i*_, and confounders 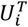. To achieve this, we substitute the above **Eq. 2** in MR with the hazard function given by **Eq. 3**, where *h*_0_(*t*) is the baseline hazard and *α*_*j*_ and *β* represent the constant (time-invariant) effects of somatic alterations and PSD on the outcome, respectively.

**Figure 1.**
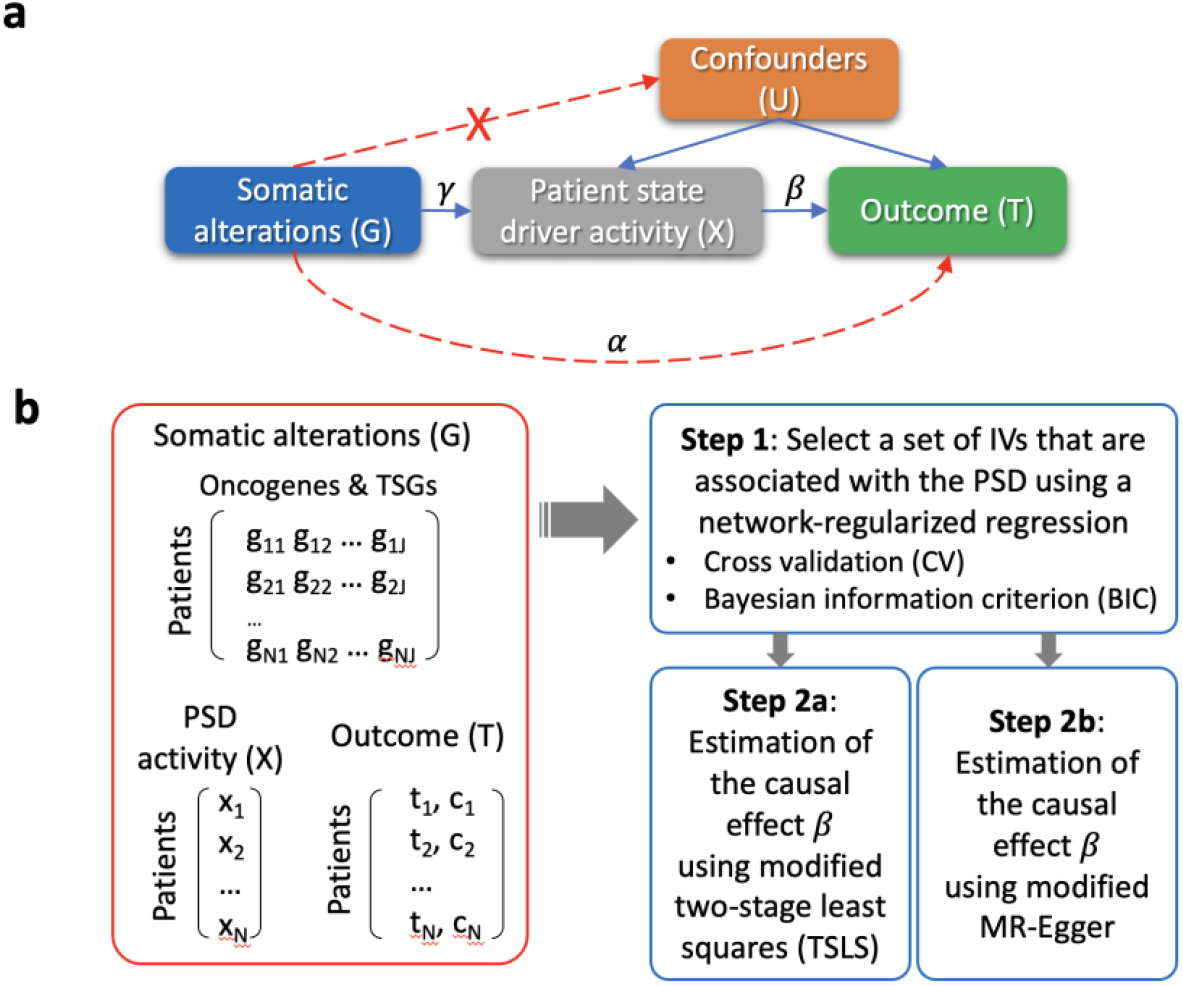
Somatic-IV workflow. (**A**) This figure illustrates the modified Mendelian Randomization (MR) assumptions for somatically acquired genetic alterations (*G*) and highlights potential violations of the assumptions with red dotted lines. The diagram shows the genetic influence on the Patient State Driver (PSD, i.e., the exposure) *X*, represented by *γ* . The direct genetic effect on the outcome *T* is denoted by *α*, while the causal effect of the PSD *X* on the outcome *T* is represented by *β*. (**B**) Somatic-IV requires three input datasets: (1) a patient-by-gene matrix *G* that represents the somatic alteration profile of a cohort, (2) an array of PSD activity *X*, and (3) censored time-to-event outcome data *T*. In the first step, Somatic-IV identifies a set of IVs associated with the PSD via a network-regularized lasso (NRL) method, employing either cross-validation (CV) or Bayesian information criterion (BIC) for selection. Subsequently, in Step 2, Somatic-IV estimates the causal effect *β* of the PSD on the outcome using either the two-stage least squares (TSLS), or the MR-Egger method.

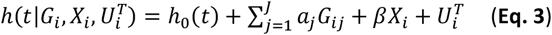

#### 2.3.1. Implementation of Somatic-IV using the two-stage regression approach

When working with individual-level data, Somatic-IV allows the estimation of the causal effect of the PSD on the outcome using a modified two-stage least squares (TSLS) method (Angrist and Imbens, 1995). The first step of this approach calculates the genetically determined component of the PSD 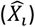, based on the sum of the effects of somatic alterations on the PSD. **Eq. 4** provides an expression to estimate 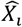, where 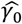 and 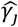 are least-squares estimates leaned from **Eq. 1**. If all somatic alterations are valid IVs (i.e., *a*_*j*_ = 0 for all variants *j*), the reduced-form **Eq. 3** can be used to estimate the causal effect of the PSD on the outcome (**Eq. 5**).

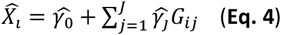

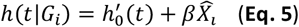

#### 2.3.2. Addressing pleiotropic effect in Somatic-IV using the MR-Egger regression

In practice, cancer somatic alterations may often violate the third IV assumption in MR (i.e., *a*_*j*_ ≠ 0). This is referred to as horizontal pleiotropy, where a genetic variant can affect an outcome directly or indirectly through pathways that are independent of the exposure being studied. To address this issue, MR-Egger regression replaces the third IV assumption with a weaker one called InSIDE (Instrument Strength Independent of Direct Effect) (Bowden, et al., 2015). MR-Egger can detect and correct for bias due to horizontal pleiotropy, leading to a bias-reduced estimation of the causal effect (Bowden, et al., 2015).

To perform MR-Egger in the Somatic-IV framework, the coefficients of each somatic alteration *j* on the outcome, denoted as 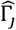, are first estimated (**Eq. 6**). A weighted regression of the genetic-outcome association estimates 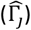 on the genetic-exposure association estimates 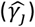 is then conducted, weighted by the inverse variance of the genetic-outcome coefficients 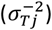 (**Eq. 7**). The intercept in Egger regression 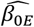 can be interpreted as an estimate of the average pleiotropic effect ( *α*_*j*_) across all the somatic alterations, while the slope 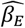 is the bias-reduced estimate of the true causal effect.

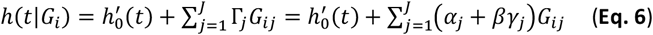

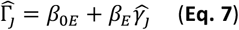

#### 2.3.3. Selecting somatic alterations using the network-regularized regression

Our Somatic-IV framework differs from previous MR or TWAS studies that use gene expression as the exposure and focus on cis-expression quantitative trait loci (cis-eQTLs) as IVs. Instead, we model trans-effects from oncogenes or tumor suppressor genes (TSGs) that carry somatic alterations to the candidate PSD of interest. Identifying somatic alterations as IVs that are both statistically and biologically meaningful is crucial for our method. To achieve this, we utilized the Reactome functional interaction (ReactomeFI) (Wu, et al., 2010) reference network, which contains information on protein interactions in biological pathways. We developed a novel method called network-regularized Lasso (NRL), which models a PSD’s protein activity as a function of the somatic alteration status of upstream oncogenes and TSGs. NRL uses an *l*1-norm to regulate model complexity and introduces a weight that penalizes oncogene-/TSG-to-PSD interactions based on their network proximity, ensuring that the regression model maximizes the biological relevance between oncogenes/TSGs and PSDs. Specifically, NRL’s optimization problem is defined in **Eq. 8**, where *p*_*j*_ represents the network proximity from the PSD under investigation to gene *j*.

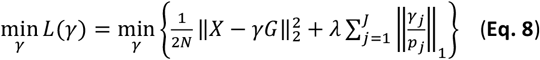

The network proximity score *p*_*j*_ is calculated from a random walk with restart (RWR) process that iteratively solves **Eq. 9** (Hofree, et al., 2013). In this equation, *P*^(0)^ is the row vector with the length of *J* genes in the network. For the gene that is the PSD under investigation, the corresponding element is 1, while for other genes, it is 0. The degree-normalized adjacency matrix of the protein interaction network is represented by *Q*. The RWR process continues until convergence is reached, which occurs when *P*^(*t*+1)^ ≈ *P*^(*t*)^. At this point, the stationary random walk score *p*_*j*_ is obtained, which represents the network proximity between the PSD and gene *j*, with *j* being the corresponding element of *P*^(*t*)^.

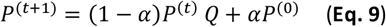

We employed two strategies to minimize the loss function *L*(*γ*) and select the optimal set of IVs: a cross-validation (CV) procedure and the Bayesian information criterion (BIC). The optimal *λ* was chosen based on the minimum CV error or BIC. We will discuss the comparison of these two approaches based on the experiments conducted on the simulated data in the following sections.

### 2.4. Generating simulated data for investigating the statistical properties of Somatic-IV framework

To assess the performance of the Somatic-IV framework, we conducted a simulation study using the R package ‘simsurv’ to generate data with parameters to reflect the distribution and characteristics of real datasets. We generated data based on the following model (**Eqs. 10** and **11**):

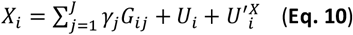

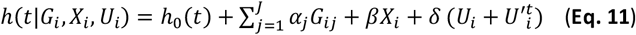

We simulated *J* = 100 genes with somatic alteration rates similar to those observed in real-world patient data, where some genes have recurrent alterations and others have rare alterations. We randomly selected 90 genes and set *γ*_*j*_ to 0, representing genetic alterations with no effect on the PSD of interest. For the remaining 10 genes, we randomly sampled IV strength *γ*_*j*_ from a Uniform(0.01, 0.75) distribution with *N* = 100, 200, 300, 500, 1000 subjects to simulate different statistical power scenarios. Error variables *U*_*i*_ , 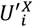 and 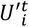 were independently generated from a *N*(0, 1) distribution. We generated α under three scenarios: (a) no pleiotropy, where *α*_*j*_ = 0 for all *j*, (b) balanced pleiotropy, where α is centered around zero:*α*_*j*_∼Uniform(−0.002, 0.002), and (c) directional pleiotropy, where *α*_*j*_∼Uniform(0, 0.002). We compared the performance of two IV selection methods, CV or BIC, and two causal effect estimation methods, TSLS or MR-Egger, under a null (*β* = 0) and positive (*β* = 0.005) causal effect as the ground truth. We computed the true positive rate (TPR) and false positive rate (FPR) at a 5% significance level.

## 3. Results

In Section 3.1, we present the results of simulations designed to investigate the statistical properties of the Somatic-IV framework under different settings. Next, in Sections 3.2 and 3.3, we demonstrate the application of this framework to identify key drivers of pancreatic adenocarcinoma (PDAC) and colorectal cancer (CRC) disease progression, respectively.

### 3.1. Somatic-IV performance on simulated data

We evaluated the performance of the Somatic-IV framework on simulated datasets by comparing two IV selection methods (CV and BIC) and two causal effect estimation methods (TSLS and MR-Egger) under different sample sizes and pleiotropic effect scenarios. The simulation study results, based on 1000 simulated datasets for each scenario, are presented in **Tables 1-3**. We implemented the TSLS and MR-Egger methods as described in the Methods section. The FPR and TPR were defined as the rejection rate of the causal null hypothesis under the null (*β* = 0) and positive (*β* = 0.005) ground truth causal effect, respectively, at a nominal significance level of 5%.

**Table 1.** Performance comparison of Somatic-IV framework using TSLS and MR-Egger estimation methods and CV and BIC IV selection methods in a simulation study under scenario (a) no pleiotropy (*α* = 0).

|  |  | TSLs |  |  |  | MR-Egger |  |  |  |
| --- | --- | --- | --- | --- | --- | --- | --- | --- | --- |
|  |  | CV |  | BIC |  | CV |  | BIC |  |
| | N | FPR | $\hat{\beta}$ (mean ± s.d.) | FPR | $\hat{\beta}$ (mean ± s.d.) | FPR | $\hat{\beta}$ (mean ± s.d.) | FPR | $\hat{\beta}$ (mean ± s.d.) |
| Ground truth<br>$\beta = 0$ | 100 | 0.21 | 0.0004 ± 0.0005 | 0.14 | 0.0005 ± 0.0022 | 0.08 | 0.0001 ± 0.0005 | 0.01 | 0.0003 ± 0.0029 |
| | 200 | 0.15 | 0.0002 ± 0.0003 | 0.07 | 0.0001 ± 0.0003 | 0.06 | -0.0001 ± 0.0003 | 0.01 | 0 ± 0.0013 |
| | 300 | 0.11 | 0.0001 ± 0.0002 | 0.06 | 0.0001 ± 0.0003 | 0.06 | -0.0001 ± 0.0002 | 0.01 | 0 ± 0.0006 |
| | 500 | 0.09 | 0.0001 ± 0.0001 | 0.05 | 0 ± 0.0002 | 0.06 | -0.0001 ± 0.0002 | 0.01 | 0 ± 0.0004 |
| | 1000 | 0.07 | 0 ± 0.0001 | 0.05 | 0 ± 0.0001 | 0.06 | -0.0001 ± 0.0001 | 0.02 | 0 ± 0.0002 |
| | N | TPR | $\hat{\beta}$ (mean ± s.d.) | TPR | $\hat{\beta}$ (mean ± s.d.) | TPR | $\hat{\beta}$ (mean ± s.d.) | TPR | $\hat{\beta}$ (mean ± s.d.) |
| Ground truth<br>$\beta = 0.005$ | 100 | 0.99 | 0.0042 ± 0.0016 | 0.91 | 0.0076 ± 0.018 | 0.79 | 0.0016 ± 0.0007 | 0.02 | 0.0047 ± 0.0098 |
| | 200 | 1.00 | 0.0036 ± 0.0007 | 0.99 | 0.0047 ± 0.0014 | 0.99 | 0.0019 ± 0.0005 | 0.23 | 0.0036 ± 0.0035 |
| | 300 | 1.00 | 0.0034 ± 0.0006 | 1.00 | 0.0041 ± 0.0009 | 1.00 | 0.0021 ± 0.0005 | 0.48 | 0.0032 ± 0.0019 |
| | 500 | 1.00 | 0.0032 ± 0.0004 | 1.00 | 0.0036 ± 0.0005 | 1.00 | 0.0022 ± 0.0004 | 0.75 | 0.0030 ± 0.0011 |
| | 1000 | 1.00 | 0.0030 ± 0.0003 | 1.00 | 0.0032 ± 0.0004 | 1.00 | 0.0023 ± 0.0003 | 0.96 | 0.0029 ± 0.0006 |

In **Table 1**, the results of scenario (a) with no pleiotropy show that all method combinations return unbiased estimates for the causal effect 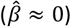 when the true causal effect is null (*β* = 0), and the FPR for the causal null hypothesis is well-controlled at close to the 5% level for all combinations of methods, especially for sample sizes of N≥300. When the true causal effect is positive (*β* = 0.005), the TPR is almost perfect (close to 1) for all method combinations, except for MR-Egger with BIC at small sample size (N≤300). However, the estimation of the causal effect 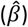 is not perfect (close to the ground truth but smaller) due to noise.

Moving on to scenario (b) with balanced pleiotropy (**Table 2**), we observe that the FPR for rejecting the causal null hypothesis is strongly inflated to 18%-54% when using the TSLS method for causal effect estimation. However, using MR-Egger regression yields better control of type 1 error rates for the causal null hypothesis. Combining MR-Egger with CV for IV selection results in a good balance of FPR (∼13%) and TPR (42%-73%). Although MR-Egger with BIC has the best control of FPR (close to 5%), it comes at the cost of reduced TPR.

**Table 2.** Performance comparison of Somatic-IV framework using TSLS and MR-Egger estimation methods and CV and BIC IV selection methods in a simulation study under scenario (b) balanced pleiotropy (*α*_*j*_∼*Unif*(−0.002, 0.002)).

|  |  | TSLs |  |  |  | MR-Egger |  |  |  |
| --- | --- | --- | --- | --- | --- | --- | --- | --- | --- |
|  |  | CV |  | BIC |  | CV |  | BIC |  |
| | N | FPR | $\hat{\beta}$ (mean ± s.d.) | FPR | $\hat{\beta}$ (mean ± s.d.) | FPR | $\hat{\beta}$ (mean ± s.d.) | FPR | $\hat{\beta}$ (mean ± s.d.) |
| Ground truth<br>$\beta = 0$ | 100 | 0.18 | 0.0001 ± 0.0005 | 0.21 | 0.0004 ± 0.0024 | 0.13 | -0.0001 ± 0.0006 | 0.01 | 0.0002 ± 0.0038 |
| | 200 | 0.26 | 0.0001 ± 0.0004 | 0.26 | 0.0001 ± 0.0006 | 0.12 | -0.0001 ± 0.0004 | 0.05 | 0 ± 0.0013 |
| | 300 | 0.30 | 0.0001 ± 0.0003 | 0.31 | 0 ± 0.0004 | 0.14 | -0.0001 ± 0.0004 | 0.08 | 0 ± 0.0009 |
| | 500 | 0.42 | 0.0000 ± 0.0003 | 0.42 | 0 ± 0.0003 | 0.14 | -0.0001 ± 0.0003 | 0.07 | 0 ± 0.0007 |
| | 1000 | 0.54 | 0 ± 0.0002 | 0.54 | 0 ± 0.0003 | 0.13 | -0.0001 ± 0.0003 | 0.08 | 0 ± 0.0005 |
| | N | TPR | $\hat{\beta}$ (mean ± s.d.) | TPR | $\hat{\beta}$ (mean ± s.d.) | TPR | $\hat{\beta}$ (mean ± s.d.) | TPR | $\hat{\beta}$ (mean ± s.d.) |
| Ground truth<br>$\beta = 0.005$ | 100 | 0.89 | 0.0023 ± 0.0018 | 0.69 | 0.0046 ± 0.0084 | 0.42 | 0.0009 ± 0.0010 | 0.01 | 0.0031 ± 0.0088 |
| | 200 | 0.97 | 0.0019 ± 0.0012 | 0.90 | 0.0025 ± 0.0019 | 0.56 | 0.0010 ± 0.0008 | 0.09 | 0.0017 ± 0.0055 |
| | 300 | 0.99 | 0.0017 ± 0.0010 | 0.96 | 0.0021 ± 0.0013 | 0.58 | 0.0010 ± 0.0008 | 0.18 | 0.0015 ± 0.0021 |
| | 500 | 1.00 | 0.0016 ± 0.0010 | 0.99 | 0.0018 ± 0.0012 | 0.62 | 0.0010 ± 0.0008 | 0.27 | 0.0014 ± 0.0016 |
| | 1000 | 1.00 | 0.0016 ± 0.0010 | 1.00 | 0.0017 ± 0.0011 | 0.73 | 0.0011 ± 0.0007 | 0.35 | 0.0013 ± 0.0013 |

Similarly, scenario (c) with directional pleiotropy (**Table 3**) shows similar results to scenario (b). Under a null causal effect (*β* = 0) as the ground truth, the FPR is strongly inflated when using the TSLS method, and it yields biased *β* estimation due to directional *α*. However, combining MR-Egger with CV achieves a good balance between FPR (6%-16%) and TPR (27%-96%). MR-Egger with BIC has the best control of FPR (close to 5%), but at the cost of reduced TPR.

**Table 3.** Performance comparison of Somatic-IV framework using TSLS and MR-Egger estimation methods and CV and BIC IV selection methods in a simulation study under scenario (c) directional pleiotropy (*α*_*j*_∼*Unif*(0, 0.002)).

|  |  | TSLs |  |  |  | MR-Egger |  |  |  |
| --- | --- | --- | --- | --- | --- | --- | --- | --- | --- |
|  |  | CV |  | BIC |  | CV |  | BIC |  |
| | N | FPR | $\hat{\beta}$ (mean ± s.d.) | FPR | $\hat{\beta}$ (mean ± s.d.) | FPR | $\hat{\beta}$ (mean ± s.d.) | FPR | $\hat{\beta}$ (mean ± s.d.) |
| Ground truth<br>$\beta = 0$ | 100 | 0.18 | 0.003 ± 0.005 | 0.14 | 0.007 ± 0.019 | 0.06 | 0.0007 ± 0.0037 | 0 | 0.0033 ± 0.0392 |
| | 200 | 0.22 | 0.003 ± 0.003 | 0.18 | 0.003 ± 0.004 | 0.07 | 0.0011 ± 0.0029 | 0.01 | 0.0005 ± 0.0141 |
| | 300 | 0.29 | 0.0023 ± 0.0020 | 0.26 | 0.003 ± 0.003 | 0.08 | 0.0013 ± 0.0022 | 0.01 | 0.0005 ± 0.0064 |
| | 500 | 0.46 | 0.0025 ± 0.0017 | 0.44 | 0.003 ± 0.002 | 0.13 | 0.0014 ± 0.0019 | 0.02 | 0.0005 ± 0.0042 |
| | 1000 | 0.64 | 0.0023 ± 0.0013 | 0.63 | 0.002 ± 0.001 | 0.16 | 0.0015 ± 0.0014 | 0.04 | 0.0003 ± 0.0028 |
| | N | TPR | $\hat{\beta}$ (mean ± s.d.) | TPR | $\hat{\beta}$ (mean ± s.d.) | TPR | $\hat{\beta}$ (mean ± s.d.) | TPR | $\hat{\beta}$ (mean ± s.d.) |
| **Ground truth**<br>**$\beta = 0.005$** | 100 | 0.69 | $0.010 \pm 0.006$ | 0.50 | $0.016 \pm 0.016$ | 0.27 | $0.0052 \pm 0.0045$ | 0 | $0.0078 \pm 0.0268$ |
| | 200 | 0.84 | $0.009 \pm 0.004$ | 0.75 | $0.012 \pm 0.006$ | 0.43 | $0.0057 \pm 0.0034$ | 0.02 | $0.0048 \pm 0.0159$ |
| | 300 | 0.96 | $0.008 \pm 0.003$ | 0.91 | $0.010 \pm 0.004$ | 0.62 | $0.0060 \pm 0.0027$ | 0.07 | $0.0053 \pm 0.0077$ |
| | 500 | 0.99 | $0.008 \pm 0.002$ | 0.99 | $0.009 \pm 0.003$ | 0.81 | $0.0063 \pm 0.0023$ | 0.17 | $0.0055 \pm 0.0049$ |
| | 1000 | 1.00 | $0.008 \pm 0.002$ | 1.00 | $0.008 \pm 0.002$ | 0.96 | $0.0065 \pm 0.0017$ | 0.36 | $0.0054 \pm 0.0033$ |

Overall, these simulations systematically benchmarked the performance of the Somatic-IV framework and informed the choice of IV selection and causal estimation methods. The best performance was achieved by combining MR-Egger with CV, and we used this combination for the study of patient datasets in the following sections.

### 3.2. Causal inference in pancreatic cancer

After conducting simulation tests to benchmark the Somatic-IV framework, we applied it to identify disease progression drivers in cancer patients. First, we focused on pancreatic ductal adenocarcinoma (PDAC), which has an extremely poor prognosis and poses a major unmet medical need. With the lowest 5-year survival rate among all cancers, at only 10%, PDAC is the third most common cause of cancer-related deaths in the United States (Siegel, et al., 2021). Our objective here was to identify PDAC patient state drivers (PSDs) that mediate the causal effect of upstream genetic alterations on patient survival.

#### 3.2.1. Pancreatic cancer data processing

We utilized data from 167 PDAC tumors from TCGA (Cancer Genome Atlas Research Network. Electronic address and Cancer Genome Atlas Research, 2017) to collect and integrate somatic mutations and CNAs in known oncogenes and tumor suppressor genes (TSGs) (Futreal, et al., 2004). We only considered likely activating alterations (missense mutations or amplification) as genetic alteration events for oncogenes, while only likely inactivating alterations (non-silent mutations and deep deletion) were considered for TSGs. From the RNA-Seq transcriptomics profiles of these TCGA PDAC patients, we inferred a PDAC-specific gene regulatory network using hARACNe (Jang, et al., 2013) that connects a pool of 4,458 candidate PSDs, including transcription factors (TFs), co-factors, and signaling molecules (Ashburner, et al., 2000), and their putative downstream target genes (regulon). The regulon expression was used as a surrogate readout to infer the protein activity of each upstream candidate PSD, yielding a more robust measurement of candidate PSD activity than using the regulator mRNA expression alone (Alvarez, et al., 2016). We used progression-free interval (PFI) as the outcome measure to analyze PDAC patient outcomes.

To ensure the validity of somatic alterations as IVs with regard to IV assumption 2 (Sections 2.1 and 2.2), we explicitly tested somatic alterations against all potential confounders present in our PDAC dataset (i.e. age, gender, race, tumor stage, and tumor grade). We excluded any somatic IV candidates that showed significant associations with these confounders (FDR < 0.25). Through this approach, we identified and removed three genes, NOTCH2 and BCL9, which were associated with increased patient age, and CCNE1, which was associated with younger age (early disease onset), from the list of candidate somatic IVs.

#### 3.2.2. Identifying key drivers of PDAC patient outcome

We employed the Somatic-IV framework to analyze 1,181 candidate PSDs, which have activities linked to somatic alterations in PDAC tumors, as determined by network regularized Lasso (NRL) regression (**Methods**, R^2^ > 0.01). We identified 99 PSDs with moderate causal effects on patient PFI (**Fig. 2a**, MR-Egger test FDR < 0.25). One notable PSD is ZBED2, previously implicated as a key transcription factor (TF) with aberrant expression in more aggressive PDAC. ZBED2 has been shown to suppress interferon response genes, subsequently repressing pancreatic epithelial cell lineage identity in PDAC, leading to increased cell motility and cancer invasion (Somerville, et al., 2020). Through NRL regression, we discovered a strong association between ZBED2 activity and somatic alterations in *KRAS, TP53*, and other oncogenes and TSGs in PDAC (**Fig. 2a**, R^2^ = 0.36). Utilizing the Somatic-IV framework, we further identified that genetic effects on ZBED2 activity are significantly correlated with genetic effects on patient outcomes (PFI), indicating a causal impact of ZBED2 on PDAC disease progression (**Fig. 2b**, MR-Egger test p = 0.005, FDR = 0.22). In summary, our findings outline a causal information flow from upstream genetic alterations to worsened patient outcomes (shorter PFI), mediated by the abnormally elevated activity of ZBED2 (**Fig. 2c**).

**Figure 2.**
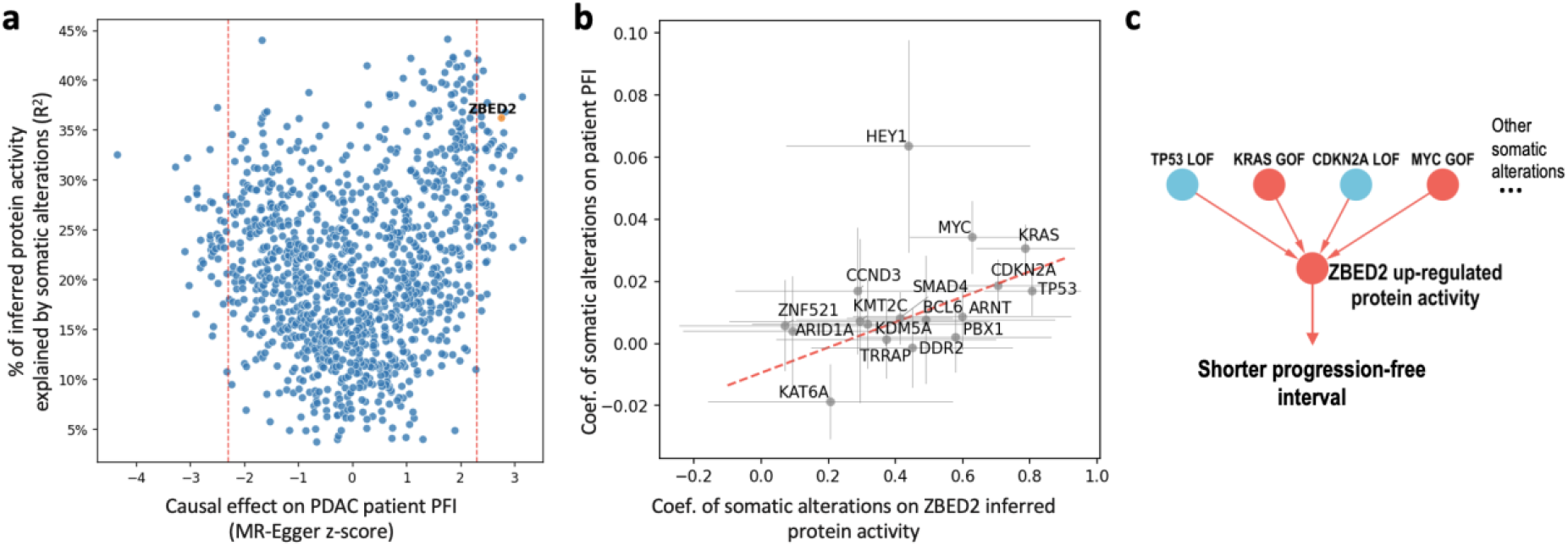
Causal inference of key drivers of PDAC patient outcome. (**A**) The graph presents the percentage of patient state driver (PSD) activity explained by somatic alterations (y-axis) plotted against the causal effect of potential PSDs on the progression-free interval (PFI) of PDAC patients (x-axis). The cutoff, denoted by red dotted lines, is set at FDR=0.25. (**B**) The plot displays the genetic-outcome (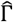 , y-axis) vs. genetic-PSD (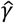 , x-axis) regression coefficients derived from a MR-Egger analysis involving 15 known cancer genes with somatic alterations linked to ZBED2 activity. The slope, indicated by a red dotted line, represents the estimated causal effect β of the PSD on the outcome, as calculated by the MR-Egger regression. (**C**) This panel presents the hypothesis that ZBED2 activity mediates the causal effect of upstream genetic alterations on patient PFI.

### 3.3. Causal inference in colorectal cancer

In our second case study, we investigated colorectal cancer (CRC). Notably, 16% of CRC cases are microsatellite instable (MSI-H), characterized by high mutation frequencies and favorable patient outcomes due to a positive response to immune-oncology therapies. This makes MSI status a likely confounder that violates IV assumption 2 (Sections 2.1 and 2.2). To avoid this violation, we specifically focused on microsatellite stable (MSS) CRC tumors, which not only represent a significant unmet medical need but also ensure the integrity of our IV assumptions. Additionally, we excluded patient tumors with a high fibrotic microenvironment (TME) to avoid bias due to tumor purity. Our aim was to identify tumor drivers that influence patient outcomes in MSS CRC cases.

#### 3.3.1. Colorectal cancer data processing

We characterized 509 colorectal cancer samples from TCGA (Cancer Genome Atlas, 2012) using the IMF classification system (Joanito, et al., 2022). From this dataset, we selected 296 non-fibrotic MSS CRC (MSS-NF CRC) tumors and integrated somatic mutations and CNAs from known oncogenes and tumor suppressor genes. Following the approach described in the PDAC case, we constructed an MSS-NF CRC-specific gene regulatory network (GRN) and inferred the activities of 4,226 candidate patient state drivers (PSDs). To analyze MSS CRC patient outcomes, we chose overall survival (OS) as our outcome measure, as it yielded more significant results compared to progression-free interval (PFI) in this context.

Similar to the PDAC case, we tested and excluded somatic IV candidates that are significantly associated with known confounders (FDR>0.25 against age, gender, race, tumor stage, and tumor grade). One gene, POLQ, was removed from the candidate somatic IVs due to its association with early onsite cancers.

#### 3.3.2. Identifying key drivers of MSS CRC patient outcome

Among the 4,226 candidate patient state drivers (PSDs), 1,024 exhibited activities linked to genetic alterations in MSS CRC tumors, as determined by network regularized Lasso (NRL) regression (**Methods**, R^2^ > 0.01). We applied the Somatic-IV framework to analyze these 1,024 candidate PSDs, identifying 193 PSDs with a statistically significant causal effect on patient overall survival (OS) (**Fig. 3a**, MR-Egger test FDR < 0.05). One PSD with the most significant causal effect on shorter overall survival time is CSNK2A2. CSNK2A2, also known as casein kinase 2 alpha prime polypeptide or CK2α’, is a catalytic subunit of protein kinase CK2. Previous studies have reported that CK2 is overexpressed in colorectal cancer tissues and can promote cell proliferation, migration, and colony formation, potentially through the phosphorylation and cytoplasm-to-nuclei translocation of β-catenin, thereby regulating Wnt signaling (Zou, et al., 2011). Employing our NRL approach, we uncovered a strong association between CSNK2A2 activity and genetic alterations in WNT signaling (*APC, AXIN2, CTNNB1, FBXW7*), RAS/PI3K signaling (*BRAF, KRAS, PTEN)*, and other oncogenes and TSGs in MSS CRC (**Figs. 3a, c**, R^2^ = 0.32). By utilizing the Somatic-IV framework, we further identified a causal effect of CSNK2A2 on MSS CRC patient OS (**Fig. 3b**, MR-Egger test p = 1.3e-07, FDR = 3.4e-05), revealing a causal information flow from upstream genetic alterations to worsened patient outcomes (shorter OS) mediated by the abnormally elevated activity of CSNK2A2 (**Fig. 3c**).

**Figure 3.**
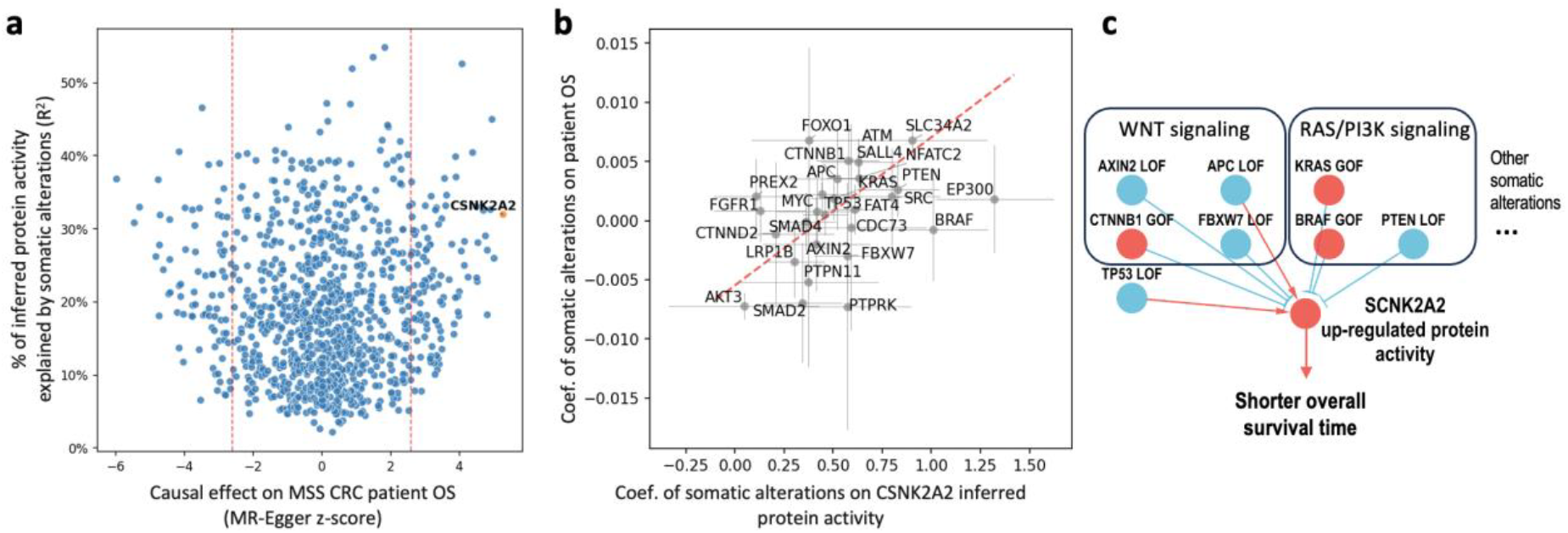
Causal inference of key drivers of MSS CRC patient outcome. (**A**) The graph presents the percentage of patient state driver (PSD) activity explained by somatic alterations (y-axis) plotted against the causal effect of potential PSDs on the overall survival (OS) of MSS CRC patients (x-axis). The cutoff, denoted by red dotted lines, is set at FDR=0.05. (**B**) The plot displays the genetic-outcome (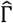 , y-axis) vs. genetic-PSD (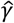 , x-axis) regression coefficients derived from a MR-Egger analysis involving 27 known cancer genes with somatic alterations linked to CSNK2A2 activity. The slope, indicated by a red dotted line, represents the estimated causal effect β of the PSD on the outcome, as calculated by the MR-Egger regression. (**C**) This panel presents the hypothesis that CSNK2A2 activity mediates the causal effect of upstream genetic alterations on patient OS.

## 4. Discussion

In this paper, we introduced the Somatic Instrumental Variable (Somatic-IV) framework, a novel approach designed to directly assess the causal effect of candidate molecular drivers on disease progression. By integrating activity measurements of a patient state driver (PSD), genetic alteration profiles of oncogenes and tumor suppressor genes (TSGs), and patient outcome data, the Somatic-IV framework can provide a bias-reduced estimation of a PSD’s causal effect on patient outcomes. Applying Mendelian Randomization (MR) in cancer studies requires specific strategies to accommodate somatic alterations (as opposed to inherited SNPs) and censored time-to-event outcome data. To address these challenges, we incorporated somatic alterations with protein interaction networks using a network-regularized Lasso (NRL) regression and performed MR in a survival context with the Aalen additive hazard model. Our case studies on pancreatic and colorectal cancer demonstrated the Somatic-IV framework’s ability to identify biologically promising candidate drivers of disease progression, highlighting its potential in generating testable target hypothesis for the development of cancer therapies.

However, it is crucial to recognize certain caveats associated with this approach, especially from a technical perspective. In the metastatic, post-standard of care (SOC) setting, treatment history could be a significant confounding factor, associated with both acquired somatic alteration and patient outcome. Cohorts with complex treatment histories requires careful patient inclusion/exclusion criteria and mathematical modeling to avoid violating the IV assumptions (Sections 2.1 and 2.2). For instance, confounding factors like targeted therapies (e.g., EGFR-TKIs) are associated with specific somatic mutations, which could violate a key IV assumption and bias the analysis. Thus, addressing these technical challenges is vital for the successful implementation of the Somatic-IV framework in real-world settings and the accurate identification of drivers of cancer progression and treatment resistance.

In conclusion, the Somatic-IV framework offers a valuable approach for identifying causal relationships between disease drivers and patient outcomes in cancer studies. Such insights could pave the way for comprehensive functional characterization, inform the development of effective therapeutic strategies, and ultimately improve patient care in the future.

